# A Reference Genome for the Critically Endangered Philippine Eagle (*Pithecophaga jefferyi*), the National Bird of the Philippines

**DOI:** 10.64898/2026.04.12.717985

**Authors:** Jonah Ray Hernandez, John Kim Aligato, Jayson Ibanez, Lorenz Rhuel Ragasa, O.P. Nicanor Austriaco

**Author notes:** These authors contributed equally to this work (Jonah Ray Hernandez, John Kim Aligato).

## Abstract

The world’s largest and rarest eagle, the Philippine Eagle (*Pithecophaga jefferyi*), also known as the monkey-eating eagle, is the national bird of the Philippines. This raptor species is endemic to the Philippine archipelago, with populations on the islands of Luzon, Leyte, Samar, and Mindanao. It is critically endangered, with an average estimated population of 392 potentially breeding pairs or 784 mature individuals. In this paper, we describe a reference genome of the Philippine Eagle (*Pithecophaga jefferyi*) from a female juvenile from the province of Nueva Ecija on the island of Luzon. We generated a *de novo* genome assembly with high contiguity and completeness, comprising 178 contigs totaling 1.345 Gbp. The genome was sequenced at a coverage of 75.2x, and Benchmarking Universal Single-Copy Orthologs (BUSCO)/Compleasm analysis yielded a BUSCO score of 99.92% (aves_odb12), corresponding to 99.7% single-copy, 0.21% duplicated, and 0.08% fragmented genes. A consensus mitogenome sequence of 19,377 bp was also generated. The genome assembly included 23,847 putative genes, and our annotation estimated that 15.78% of the genome consisted of repetitive elements. Genome heterozygosity (H) was estimated to be 0.020%, in comparison to other birds with genome heterozygosity values ranging from 0.0103% to 0.923%. Whole-genome comparisons with publicly available genomes suggest that the Philippine eagle belongs to the snake-eagle subfamily (*Circaetinae*) rather than the harpy-eagle subfamily (*Harpiinae*). Pairwise sequentially Markovian coalescent (PSMC) analysis suggests that the effective population size was around 4,000 individuals from about 100 KYA to about 1 KYA. Finally, we constructed a minimum spanning network, which revealed that our mitogenome from the northern island of Luzon occupies a peripheral position, separated from the dominant haplotype cluster found in the southern island of Mindanao by multiple mutational steps, indicating substantial mitochondrial divergence.

## INTRODUCTION

Of the approximately 10,500 known species of birds, about 225 species, which is roughly 2% of the total, are characterized as critically endangered according to the International Union for Conservation of Nature (IUCN) criteria (Maclean, 2023; p. 106). According to these IUCN criteria, a bird species is considered critically endangered if it fulfills one of four conditions: The bird population 1) has experienced a reduction in numbers of at least 80% over the past decade or three generations, whichever is longer; 2) has a geographic range of less than 100 km^2^ that is threatened in different ways; 3) is small (≤ 250 mature individuals) and in decline; and 4) has a high probability (≥50%) of going extinct in the wild within a decade or three generations, whichever is longer (IUCN Standards and Petitions Committee, 2024).

The world’s largest and rarest eagle, the Philippine Eagle (*Pithecophaga jefferyi*), also known as the monkey-eating eagle, is the national bird of the Philippines (Ramos, 1995). It is depicted on the newly released Philippine polymer one-thousand-peso banknote, and is prominently featured in the Philippine passport. This raptor species is endemic to the Philippines, with populations on the islands of Luzon, Leyte, Samar, and Mindanao (Fisher and Hicks, 2000). There is evidence that birds from Luzon possess adaptive traits that distinguish them from their counterparts on other islands of the archipelago (Abano et al., 2016). The Philippine eagle is critically endangered with an estimated average population of 392 potentially breeding pairs or 784 mature individuals (BirdLife International, 2025; DENR, 2019; Sutton et al., 2023). Significantly, the Philippine eagle ranks tenth in the world’s top 100 list of Evolutionarily Distinct and Globally Endangered (EDGE) bird species (Zoological Society of London, 2024).

Several factors contribute to the ongoing decline in the Philippine eagle population. First, the species’s biological and ecological traits make it very sensitive to human interference (Ibanez et al., 2016). The bird is a slow breeder—a monogamous eagle pair rears only one chick every two years—and it is late-maturing—a hatchling takes at least six years to become sexually mature (Ibanez, 2008). Eagle pairs also demonstrate high mate and nest site fidelity (Miranda et al., 2000). Thus, when the eagle population declines due to excessive human activity, such as shooting and trapping, it is difficult for the birds to recover on their own. Second, the Philippines has experienced significant deforestation and continues to do so, with an estimated loss of about 50,000 hectares between 2000 and 2012 (Fallarcuna and Perez, 2016). This has dramatically reduced the hunting ranges, shelters, and breeding sites available to eagles. Finally, conservation efforts, including enforcement of the 2001 Philippine Wildlife Act, to protect eagles and their habitats from human destruction remain lacking (Ibanez et al., 2016). Nonetheless, the conservation of the Philippine Eagle remains a top priority of the national and local governments of the Philippines (Araña, 2024).

Given the urgent need to increase the population of Philippine eagles, the Philippine Eagle Foundation (PEF), a private, non-profit organization in the Philippines, has developed a breeding program to raise eagles to repopulate vacant habitats in the wild and save the species from extinction (Collar and Butchart, 2013). Since its founding in 1987, PEF has produced thirty captive-bred eagles, three of which have been released into the wild (Bastian et al. 2007). Their conservation efforts would benefit from genetic information to improve breeding outcomes among captive eagles.

Convincing arguments have been made that increasing and maintaining genome-wide genetic variation is crucial for conservation work (Kardos et al., 2021; Jeon et al., 2024). A recent study by Bacus et al. (2025) of partially sequenced mitochondrial genomes from twenty-seven birds housed at the Philippine Eagle Sanctuary established by PEF revealed a mean nucleotide diversity (π) for the mitochondrial genome of 0.054%, which is significantly lower than an earlier reported π value for mtDNA from Philippine eagles of 0.62% calculated using the mitochondrial control region (Luczon et al., 2014). Genome-wide data from *de novo* whole-genome sequencing (WGS) of the Philippine eagle should help increase the robustness and reliability of measurements of heterozygosity and other calculated indices of genome diversity for this critically endangered raptor. This would give us a benchmark to evaluate efforts to conserve the species.

In this paper, we describe a reference genome for the Philippine Eagle (*Pithecophaga jefferyi*) from a female juvenile named “Binuang” from the province of Nueva Ecija on the island of Luzon. We obtained a genome assembly with high contiguity and completeness, comprising 178 contigs totaling 1.345 Gbp. The genome was sequenced at a coverage of 75.2x, and Benchmarking Universal Single-Copy Orthologs (BUSCO) analysis with Compleasm identified 99.92% complete BUSCOs (aves_odb12), corresponding to 99.7% single-copy, 0.21% duplicated, and 0.08% fragmented genes. The genome assembly included 23,847 putative genes. Genome heterozygosity (H) was estimated to be 0.020%, in comparison to other birds with genome heterozygosity values ranging from 0.0103% to 0.923% (Brüniche-Olsen et al., 2021). Next, whole-genome comparisons with publicly available genomes suggest that the Philippine Eagle belongs to the snake-eagle subfamily (Circaetinae) rather than to the harpy-eagle subfamily (Harpiinae) or the hawk subfamily (Accipitrinae). Pairwise sequentially Markovian coalescent (PSMC) analysis suggests that the effective population size was around 4,000 individuals from about 100 KYA to about 1 KYA. Finally, we constructed a minimum spanning network, which revealed that our mitogenome from the northern island of Luzon occupies a peripheral position, separated from the dominant haplotype cluster found in the southern island of Mindanao by multiple mutational steps, indicating substantial mitochondrial divergence.

## MATERIALS & METHODS

### Sample Collection

Following standard procedures, whole blood was collected on March 14, 2025, by licensed veterinarians during health assessments of a female eagle monitored in Gabaldon, Nueva Ecija, in the Philippines. The juvenile, named “Binuang”, was approximately 11 months of age at the time of capture and weighed 4.4 kg. Blood samples were placed in EDTA-treated collection tubes and transported to the lab for short-term storage at −20°C. Samples were mixed with Zymo DNA/RNA Shield in preparation for transport to Plasmidsaurus.com for DNA extraction and sequencing. Trapping, tagging, and blood sample collection were conducted under Gratuitous Permit No. III-2025-007 issued by the Department of Environment and Natural Resources (DENR) on February 13, 2025.

### DNA Extraction and Sequencing

Genomic DNA was extracted from preserved whole blood using NEB’s Monarch extraction kit (T3010L). Library preparation was performed with Oxford Nanopore’s Rapid Barcoding Kit (SQK-RBK114.96). Sequencing was performed with a PromethION P24 using an R10.4.1 flow cell, with a minimum Qscore of 10. Finally, sequencing reads were basecalled with Dorado in super-accurate mode.

### Nanopore long read processing

The original Fastq file was divided into four smaller subsets using Seqkit split2 v2.10.0 (Shen et al. 2024) to accommodate computational limitations. Each subset was then passed through Porechop v0.2.4 (Wick, 2018) to remove adapter sequences and split chimeric reads. The trimmed reads were then concatenated and filtered using Filtlong v.0.2.1 (Wick, 2018) with the parameters --keep_percent 95 & --mean_q_weight 10. Read quality metrics of both raw and processed reads were evaluated using NanoComp v1.25.3 (De Coster & Rademakers, 2023).

### Genome Assembly and Polishing

Initial assemblies were generated using Raven v1.8.3 (Vaser et al. 2021), Goldrush v1.2.2 (Wong et al. 2023), and Nextdenovo v2.5.2 (Hu et al. 2024) following the strategy described in McGrath et al. (2025). Hifiasm version v0.25.0 in its ONT mode was also used to generate the initial assembly. The resulting assemblies, together with a Flye v2.9.1 (Kolmogorov et al. 2019) assembly provided by Plasmidsaurus, were assessed using Compleasm v0.2.7 (Huang and Li 2023) across the aves odb12 database. The N50 of the different assemblies was also calculated and compared using QUAST v5.3.0 (Tesler et al. 2013). Among the initial assemblies, Hifiasm produced the most complete and contiguous result (Table 1) and was therefore polished with a single round of Medaka v2.0.1 (github.com/nanoporetech/medaka) using the r1041_e82_400bps_sup_v4.3.0 model. The resulting polished assembly was once again assessed using Compleasm and QUAST. Because the number of duplicated genes was low (0.18%), haplotig purging was deemed unnecessary to avoid losing some true contigs.

**TABLE 1:** PhEagle Assembly Features Extracted from Different De Novo Long-Read Genome Assemblers.

| ASSEMBLY | CONTIGS | TOTAL LENGTH | N50 | L50 | Compleasm/BUSCO<br>(aves_odb12) |
| --- | --- | --- | --- | --- | --- |
| ASM2572802v1(previous) | 11,238 | 1,257,274,458 | 38,333,288 | 13 | 89.38% |
| eagle1_goldrush | 5,402 | 1,203,166,060 | 7,918,162 | 37 | 95.45% |
| eagle1_flye | 2,675 | 1,284,745,667 | 22,263,027 | 19 | 74.11% |
| eagle1_raven | 810 | 1,238,446,163 | 16,444,478 | 24 | 99.41% |
| eagle1_nd | 191 | 1,246,315,321 | 28,053,043 | 15 | 99.89% |
| eagle1_hifiasm | 178 | 1,345,556,334 | 45,405,676 | 11 | 99.92% |
| eagle1_hifiasm_medaka | 178 | 1,345,718,307 | 45,411,332 | 11 | 99.92% |

### Contamination Detection

The final polished assembly was then mapped back to the initial reads using Minimap2 v2.29 (Li, 2021) and subsequently converted into a fastq file using Samtools v1.21 (Li et al. 2009). The taxon assignment of the final contigs as well as the unmapped reads was determined by Kraken2 v2.17.1 (Wood et al. 2019) on a custom database composed of Accipitridae genomes and the standard database.

### Mitogenome Assembly and Alignment

Raw reads were then mapped to the complete mitogenome of a Philippine eagle with the accession number CM137358.1, using Minimap2 v2.29 (Li, 2021). The mitochondrial reads were then sorted, indexed, and converted to fastq using Samtools v1.21 (Li et al. 2009). Variants were then identified and collapsed into one consensus sequence using Bcftools v1.22 (Li et al., 2009). The resulting circular mitogenome was then annotated using MitoZ (Meng et al. 2019). The generated mitogenome was concatenated with the reference mitogenome and previously published partial mitogenomes from Bacus et al. (2025). All mitochondrial sequences were aligned using MAFFT (Katoh & Standley 2013) with 1000 iterations on global pairing mode. The alignment gaps from both ends were then removed in MEGA12 (Kumar et al., 2024), and the file was converted to NEXUS format. The locational traits were assigned according to the three principal islands from which the samples were collected. The NEXUS file was then loaded into PopART (Leigh & Bryant 2015) to create the minimum spanning network.

### Synteny analysis

To determine synteny, the draft assembly was aligned to the Harpy eagle genome (GCF_026419915.1) using D-genies (Cabanettes and Klopp, 2018) with Minimap2 v2.28 (Li, 2021). The obtained PAF file was filtered to include only contigs with a total alignment length of at least 100 kb total alignment length. Synteny was visualized in RStudio using the RIdeogram package (Hao et al., 2020).

### Repeat Identification and Genome Annotation

Prior to genome annotation, the assembly was screened for repetitive elements using RepeatModeler (v2.0.5) (Flynn et al., 2020) to identify *de novo* repeat families. Subsequently, these identified repeats were used to soft-mask the genome assembly using RepeatMasker (v4.1.5) (Tarailo-Graovac & Chen, 2009). This step was performed to compute transposable element statistics and mask transposable elements and low-complexity regions while preserving the underlying genomic sequence for downstream gene prediction. Transfer RNA (tRNA) genes were identified using tRNAscan-SE (v2.0.12) (Chan et al., 2021) executed in eukaryotic mode (-E) utilizing Infernal-enabled covariance models. To establish a high-confidence tRNA repertoire, predicted loci were filtered based on secondary structure scores. Specifically, only those with bit scores ≥50 (or ≥60 for specialized isotypes) were retained, while all sequences annotated as pseudogenes were explicitly excluded from the final count. Additionally, the presence and total count of ribosomal RNA (rRNA) genes were determined using barrnap (v0.9) (Seemann, 2018) with the default eukaryotic kingdom setting.

Gene prediction was conducted using the BRAKER3 (v3.0.8) pipeline (Gabriel et al., 2024). In the absence of species-specific RNA-seq transcriptomic data, the pipeline was executed in protein-mode. We utilized the OrthoDB v11 vertebrata protein database (Kuznetsov et al., 2022) to provide high-quality homology evidence for training the underlying GeneMark and Augustus algorithms. To ensure high-quality gene models, the initial predictions were filtered to remove transcripts shorter than 50 amino acids or those containing premature stop codons. All remaining BRAKER3 parameters were left at their default settings. The completeness of the final gene set was assessed using BUSCO (v5.5.0) (Simão et al., 2015) against the aves_odb12 lineage to ensure the recovery of highly conserved orthologs.

### Population Dynamics

To preserve allelic diversity for demographic modeling, we generated a diploid consensus sequence by integrating filtered variants into the reference assembly using BCFtools (v1.22) consensus (Li et al., 2009). We employed the —iupac-codes flag to incorporate IUPAC ambiguity symbols at heterozygous sites (e.g., ‘R’ for A/G transitions), thereby maintaining the diploid state of the individual and avoiding the loss of information associated with haploid collapsing. Historical effective population size (*Ne*) was reconstructed using the Pairwise Sequentially Markovian Coalescent (PSMC) v0.6.5 (Li & Durbin, 2011). To scale the results to absolute time and population size, we utilized a mean generation time (g) and a mutation rate (μ). Following the methodology of Vilaça et al. (2021), g was calculated as age at maturity+(0.5×reproductive longevity). Based on life-history data for the Philippine Eagle—specifically an age to maturity of 6 years and a reproductive longevity of 25 years—we set a generation time of 18.5 years. The demographic trajectories were scaled using a mutation rate of 1.91 x 10^-9^ substitutions per site per year (Li et al., 2009). The PSMC analysis was executed with the following parameters: -N25 -t15 -r5 -p “4+25*2+4+6”. To assess the statistical robustness of the inferred demographic fluctuations and estimate confidence intervals, 100 bootstrap replicates were performed by randomly resampling the original genome segments (Li & Durbin, 2011).

### Phylogenetic Analysis

To reconstruct the species tree, peptides from annotated genome datasets for 13 species were retrieved from NCBI Datasets. This dataset comprises eight members of the Accipitridae family, with representatives from Cathartidae, Pandionidae, and Sagittariidae serving as outgroups. To increase taxon sampling, peptide sequences were also extracted from six additional unannotated Accipitridae genomes by aligning the eight annotated Accipitridae proteomes to these genomes using Miniprot v0.18 (Li, 2023). The resulting peptide sets were then used as input for OrthoFinder v3.1.0 (Emms et al., 2025) to identify orthologs under default parameters.

## RESULTS

### Assembly and Annotation

We have generated a near-complete genome assembly for the Philippine Eagle (*Pithecophaga jefferyi*), which we have named UST_PhEagle 1.0 (Table 1). We obtained 101.2 Gb of Oxford Nanopore reads with a coverage of 75.2X (Table 2 & Figure 1). The final curated assembly comprised 1,345,718,307 bases across 178 contigs, with an N50 of 45.4 Mb. Of these, eleven are full chromosomes with intact telomere ends (Figure 2B). Benchmarking Universal Single-Copy Orthologs (BUSCO)/Compleasm analysis yielded a BUSCO score of 99.92% (aves_odb12), corresponding to 99.7% single-copy, 0.21% duplicated, 0.08% fragmented, and 0.02% missing genes. The repetitive content was initially estimated from sequencing reads at 9.14% of the total genome length. We identified 269,749 single-nucleotide polymorphisms (SNPs) and calculated heterozygosity at 0.020% (Table 3). The UST_PhEagle 1.0 reference genome is deposited under the assembly accession number XXX at NCBI.

**FIGURE 1:**
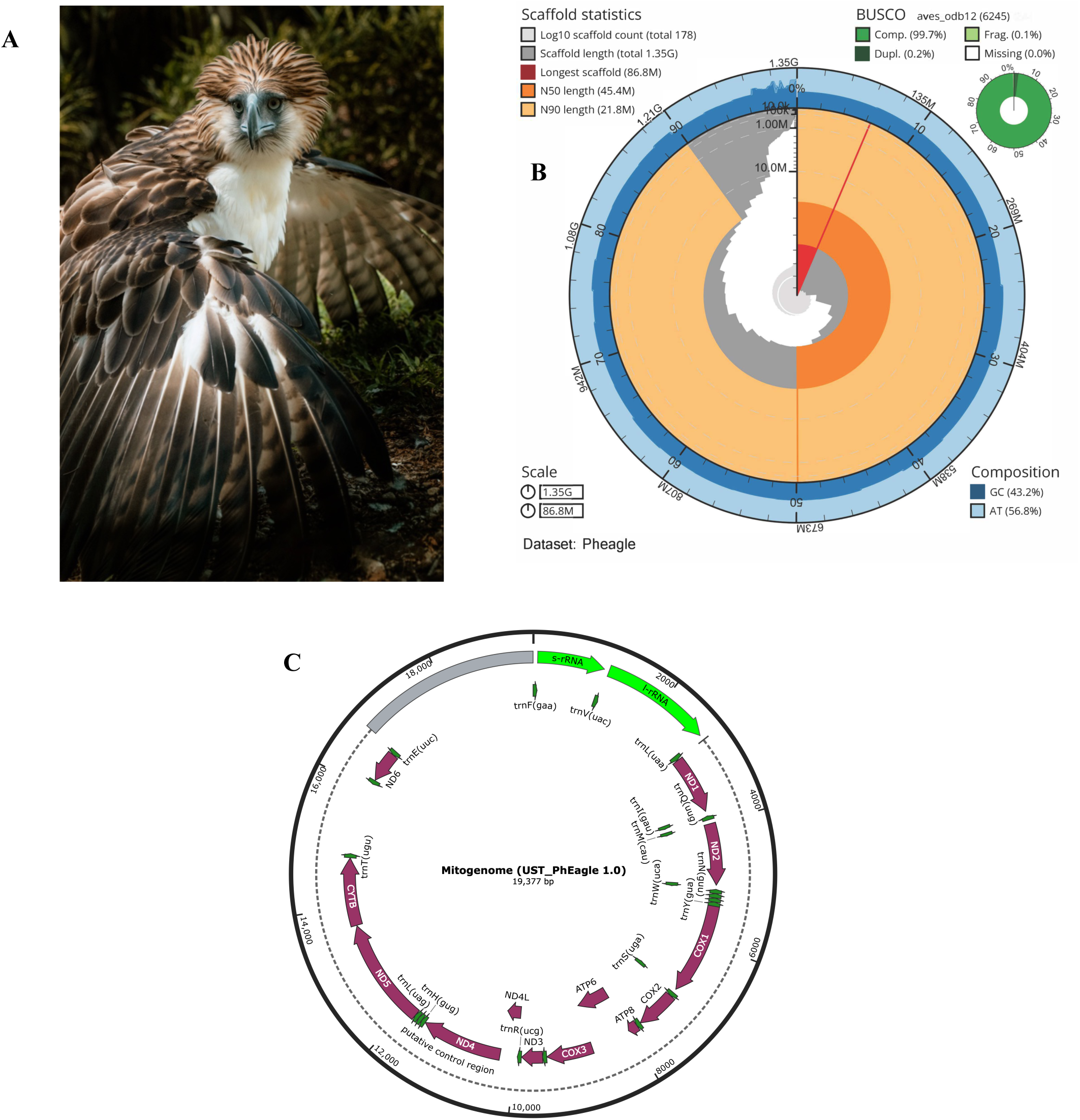
Genome Assembly Statistics and the Mitogenome for the Philippine Eagle (*Pithecophaga jefferyi*) Genome Assembly, UST_PhEagle 1.0A

**FIGURE 2:**
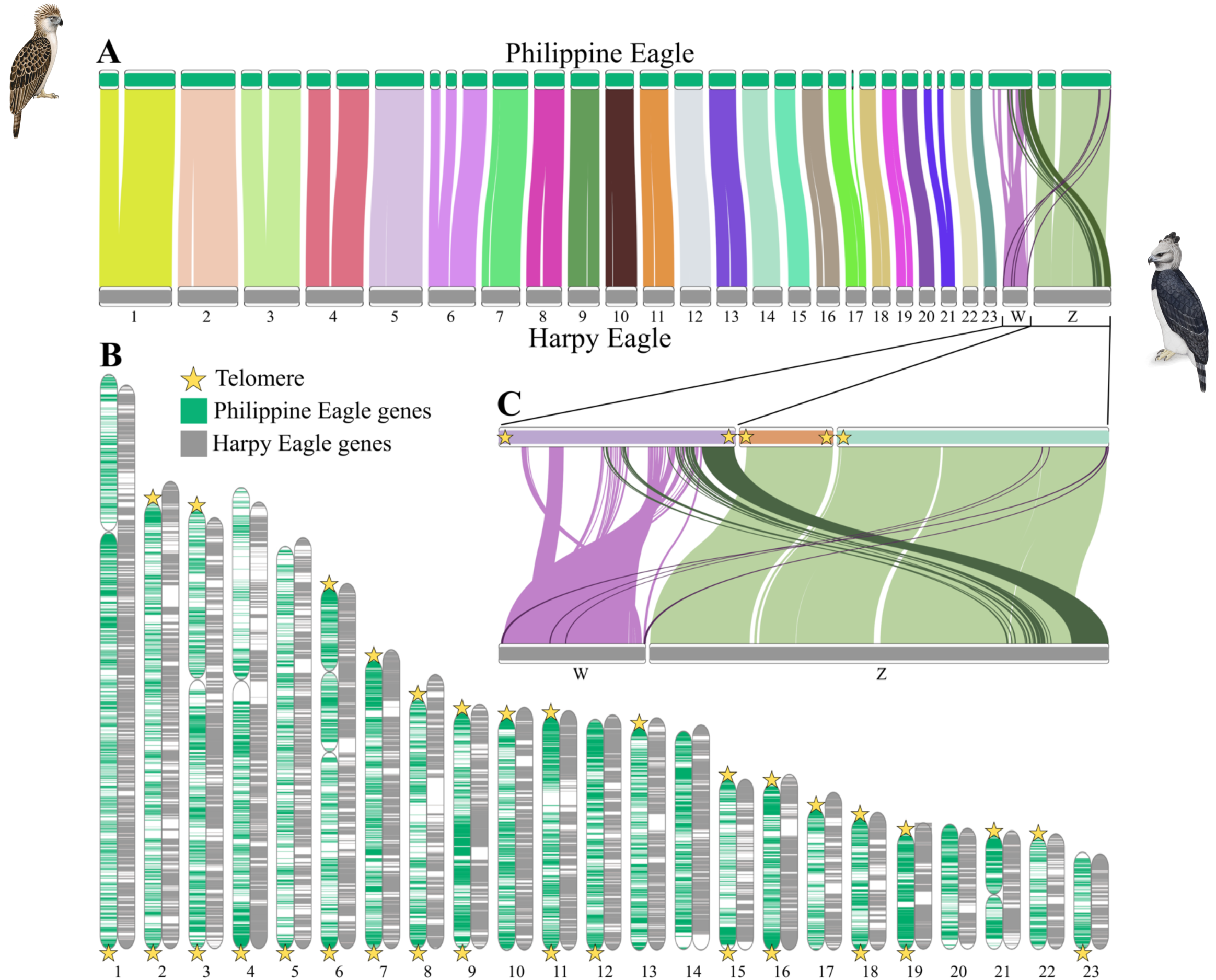
Whole-Genome Synteny Plot Comparing the Philippine Eagle (*Pithecophaga jefferyi*) and Harpy Eagle (*Harpia harpyja*)

**TABLE 2:** Summary Statistics for the Philippine Eagle (*Pithecophaga jefferyi*) Genome Assembly, UST_PhEagle 1.0.

|  | UST_PhEagle 1.0 |
| --- | --- |
| Total Length | 1,345,718,307 bp |
| Contigs | 178 |
| GC (%) | 43.19% |
| N50 | 45,411,332 |
| L50 | 4 |
| Depth of Sequencing | 75.2x |
| #Ns per 100 Kbp | 0 |
| Compleasm/BUSCO (aves_odb12) | 99.92% |
| Single Copy | 99.7% |
| Duplicated | 0.21% |
| Fragmented | 0.08% |
| Missing | 0.02% |
| Single Copy Genes (aves_odb12) | 6,245 |
| Mitogenome | 19,377 bp |

**TABLE 3:** Comparative Statistics for Heterozygosity Conservation Status: LC (Least Concern); VU (Vulnerable); EN (Endangered); CR (Critically Endangered)

|  | Conservation Status | Heterozygosity (H) |
| --- | --- | --- |
| Buderigar<br>(*Melopsittacus undulatus*) | LC | 0.431%<br>(Li et al., 2014) |
| Striped Tit-Babbler<br>(*Mixornis gularis*) | LC | 0.23% to 0.26%<br>(Tan et al., 2018) |
| Chicken<br>(*Gallus gallus*) | LC | 0.10% to 0.30%<br>(Xu et al., 2023) |
| Little Egret<br>(*Egretta garzetta*) | LC | 0.251%<br>(Li et al., 2014) |
| Bald Eagle<br>(*Haliaeetus leucocephalus*) | LC<br>(Once EN) | 0.043%<br>(Li et al., 2014) |
| White-tailed Eagle<br>(*Haliaeetus albicilla*) | LC<br>(Once VU) | 0.04%<br>(Li et al., 2014) |
| Crested Ibis<br>(*Nipponia nippon*) | EN | 0.043%<br>(Li et al., 2014) |
| Philippine Eagle<br>(*Pithecophaga jefferyi*) | CR | 0.020%<br>(this study) |
| Average Heterozygosity<br>(32 LC Species) | LC | 0.249%<br>(Li et al., 2014) |
| Average Heterozygosity<br>(8 EN-VU Species) | EN-VU | 0.0108%<br>(Li et al., 2014) |

Using a complete Philippine eagle mitogenome as a reference (GenBank CM137358.1), we obtained 1,875 mitochondrial reads with 269X coverage across the full mitogenome. A consensus mitogenome sequence of 19,377 bp was also generated. MitoZ annotation revealed that the consensus mitogenome sequence is circular and contains the complete set of 13 protein-coding genes, 22 tRNA genes, and 2 rRNA genes (Figure 1C).

The BRAKER3 annotation pipeline annotated 23,847 unique protein-coding loci, spanning 26,467 mRNAs with 167,868 exons and 141,430 introns, 331 tRNAs, and 853 rRNAs (Table 4). We compared the annotated Philippine Eagle (*Pithecophaga jefferyi*) coding sequences with those of four other avian species: the Eurasian Goshawk (*Accipiter gentilis*), the Golden Eagle (*Aquila chrysaetos*), the California Condor (*Gymnogyps californianus*), and the Chicken (*Gallus gallus*). Orthology inference using OrthoFinder assigned 195,835 genes (91.1%) to 20,982 orthogroups, indicating high annotation completeness and consistency. A total of 19,210 genes (8.9%) remained unassigned, consistent with expected levels of lineage-specific or rapidly evolving sequences. We identified 1,694 species-specific orthogroups across all five analyzed avian species, comprising 8,054 genes (3.7%), suggesting moderate lineage-specific gene innovation without evidence of annotation inflation. A total of 259 orthogroups comprising 749 genes were identified as unique to the Philippine eagle, representing approximately 3% of the annotated gene set. This relatively low level of lineage-specific gene content is consistent with the conserved genomic architecture observed across avian species and suggests that species-specific adaptations are more likely driven by regulatory evolution or gene family expansions rather than the emergence of novel genes.

**TABLE 4:** Summary Statistics for the Philippine Eagle (*Pithecophaga jefferyi*) BRAKER3 Genome Annotation.

|  | UST_PhEagle 1.0<br>(BRAKER3) |
| --- | --- |
| # of Coding Sequences/Genes | 23,847 |
| # of mRNAs | 26,467 |
| Exons | 167,868 |
| Introns | 141,430 |
| tRNAs | 331 |
| rRNAs | 853 |
| % of TE<br>(Transposable Elements) | 15.78% |
| Retroelements | 177,396 (7.37%) |
| DNA Transposons | 13,509 (0.13%) |
| Uncharacterized Elements | 83,837 (7.05%) |

Finally, the Philippine eagle genome exhibits a moderately elevated transposable element (TE) content (∼15.8%), placing it at the upper range of avian genomes (Table 4). The TE landscape is strongly dominated by retroelements, consistent with the canonical prevalence of CR1-like LINEs in birds, while DNA transposons are nearly absent (Galbraith et al., 2021). Notably, a substantial fraction of repeats (∼7%) remains uncharacterized, likely representing highly diverged or lineage-specific elements.

### Comparative Genomics and Phylogenetics

We found strong synteny between the genomes of our Philippine Eagle (*Pithecophaga jefferyi*) and the Harpy Eagle (*Harpia harypyja*), which has one of the best reference genomes of the *Accipitriformes* (Canesin et al., 2024), preserving numerous collinear gene blocks (Figure 2). The largest autosomal macrochromosomes exhibit clean, uninterrupted syntenic blocks with few crossing lines and little evidence of major translocations, indicating strong conservation of karyotypic structure within Accipitridae (Figure 2B). In contrast, the sex chromosomes, W and Z, reveal complex rearrangement signals, suggesting intra-chromosomal inversions (Figure 2C). The Z chromosome retained relatively large, contiguous syntenic blocks, whereas the W chromosome showed fragmented and highly rearranged synteny, with multiple discontinuous alignments to the Z chromosome. This pattern is consistent with the well-documented degeneration and repeat-driven restructuring of the avian W chromosome following recombination suppression (Sigeman et al., 2021; Sigeman et al., 2024). W chromosomes are also notoriously difficult to assemble due to their repeat-rich, heterochromatic nature.

To determine the phylogenetic relationships among the Philippine eagle and select members of Accipitridae, we generated a phylogenetic tree with sequences from Cathartidae, Sagittariidae, and Pandionidae as outgroups (Figure 3). Our analysis recovered a well-resolved topology placing *Pithecophaga jefferyi* within the serpent-eagle radiation, where it formed a sister relationship with *Spilornis perplexus* and was closely associated with *Circaetus pectoralis.* This assemblage was distinct from other major accipitrid lineages, including the accipitrine hawks, the buteonine buzzards, the sea eagles, and the aquiline eagles.

**FIGURE 3:**
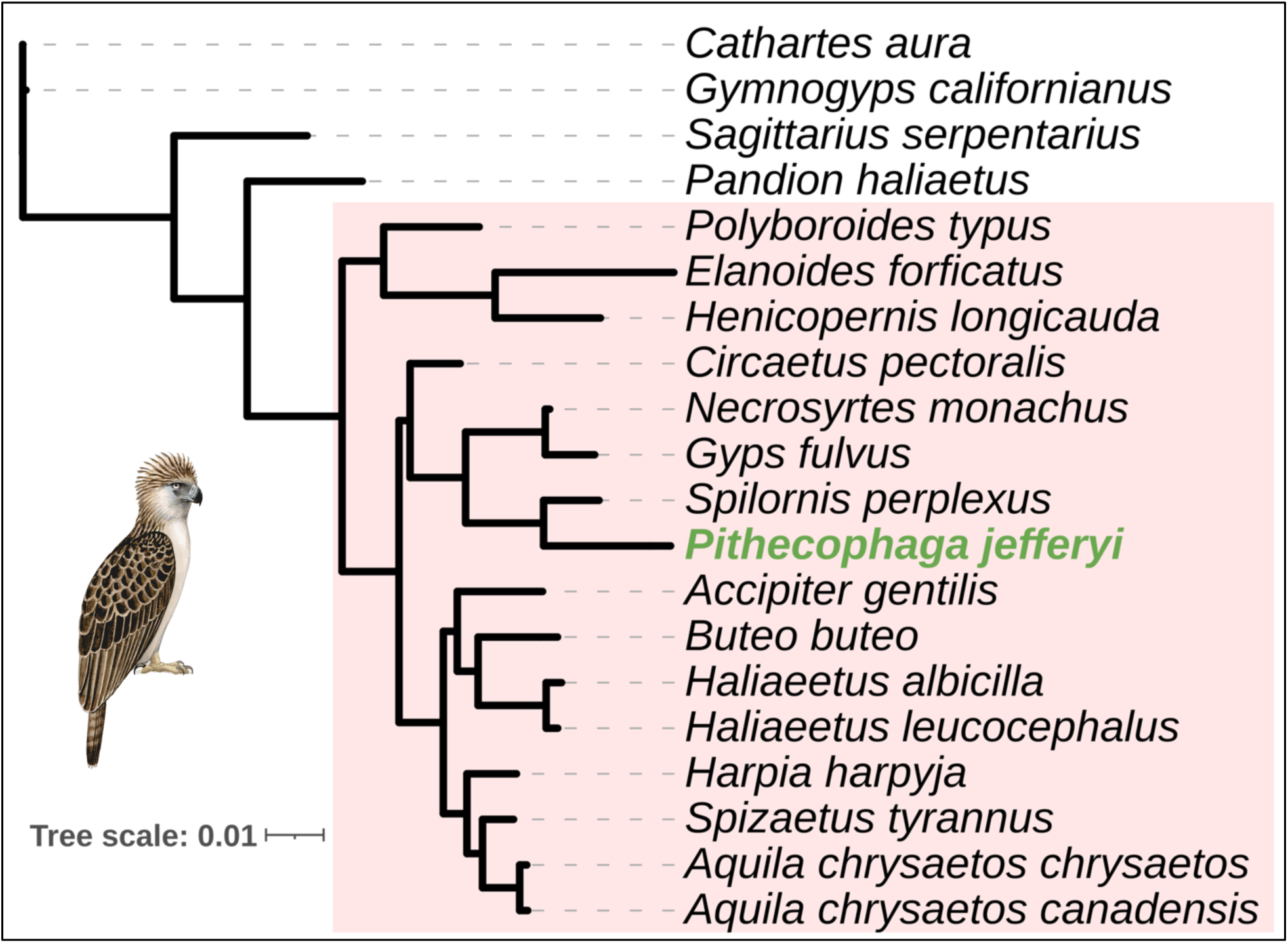
Phylogenetic Tree of *Pithecophaga jefferyi*

### Past Effective Population Size Dynamics

We reconstructed the past demographic history of the Philippine eagle using our *de novo* assembly (Figure 4). The pairwise sequentially Markovian coalescent (PSMC) analysis suggests a prominent population expansion peak of about 45,000 individuals during the Mid-Pleistocene (∼100 KYA to 1 MYA), followed by a steep decline. This decline reached an effective population size of around 4,000 individuals about 100,000 YA, which remained stable through the Last Glacial Maximum (18-23 KYA) and most of the Holocene (12-1 KYA), two sequential periods characterized by opposing cooling and warming phases, respectively.

**FIGURE 4:**
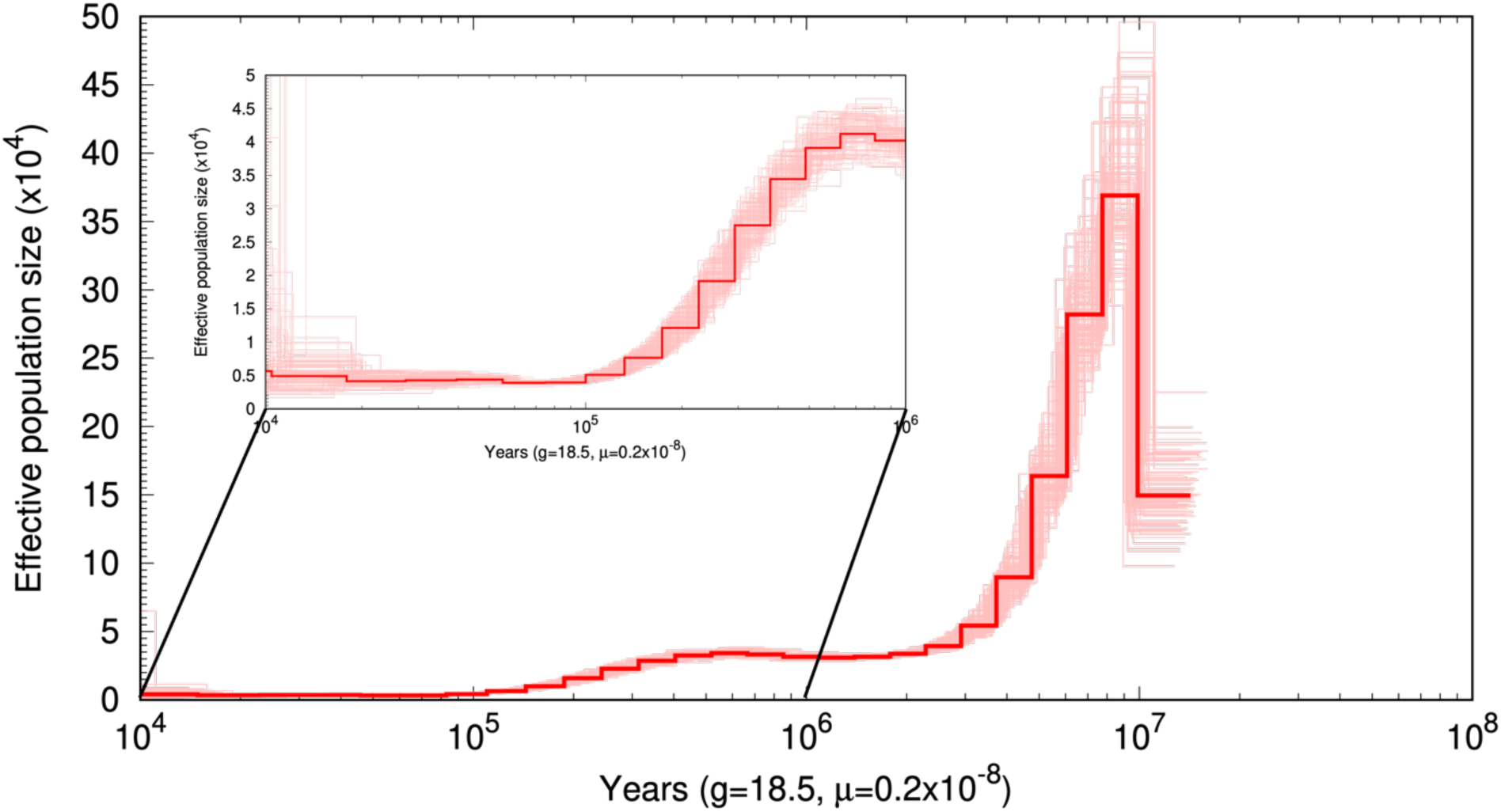
Past Effective Population Size (*N_e_*) estimates of the Philippine Eagle. The darker red line depicts the *N_e_* through time, while lighter colors represent bootstrap estimates over 100 replicates.

### Mitochondrial Haplotype Network Analysis

Using our Philippine Eagle mitogenome, we constructed a minimum spanning network to determine the genetic relationships of our eagle, “Binuang,” with 32 other eagles whose partial mitogenomes had been recently described (Bacus et al., 2025). Our mitogenome from the island of Luzon occupies a peripheral position, separated from the dominant haplotype cluster on the island of Mindanao by multiple mutational steps, indicating substantial mitochondrial divergence (Figure 5). This pattern is consistent with either historical geographic isolation or the persistence of a distinct maternal lineage, potentially reflecting population structuring across the three major island groups of the archipelago. The absence of closely related haplotypes further supports the interpretation that our Luzon eagle represents a relatively rare or undersampled lineage within the species.

**FIGURE 5:**
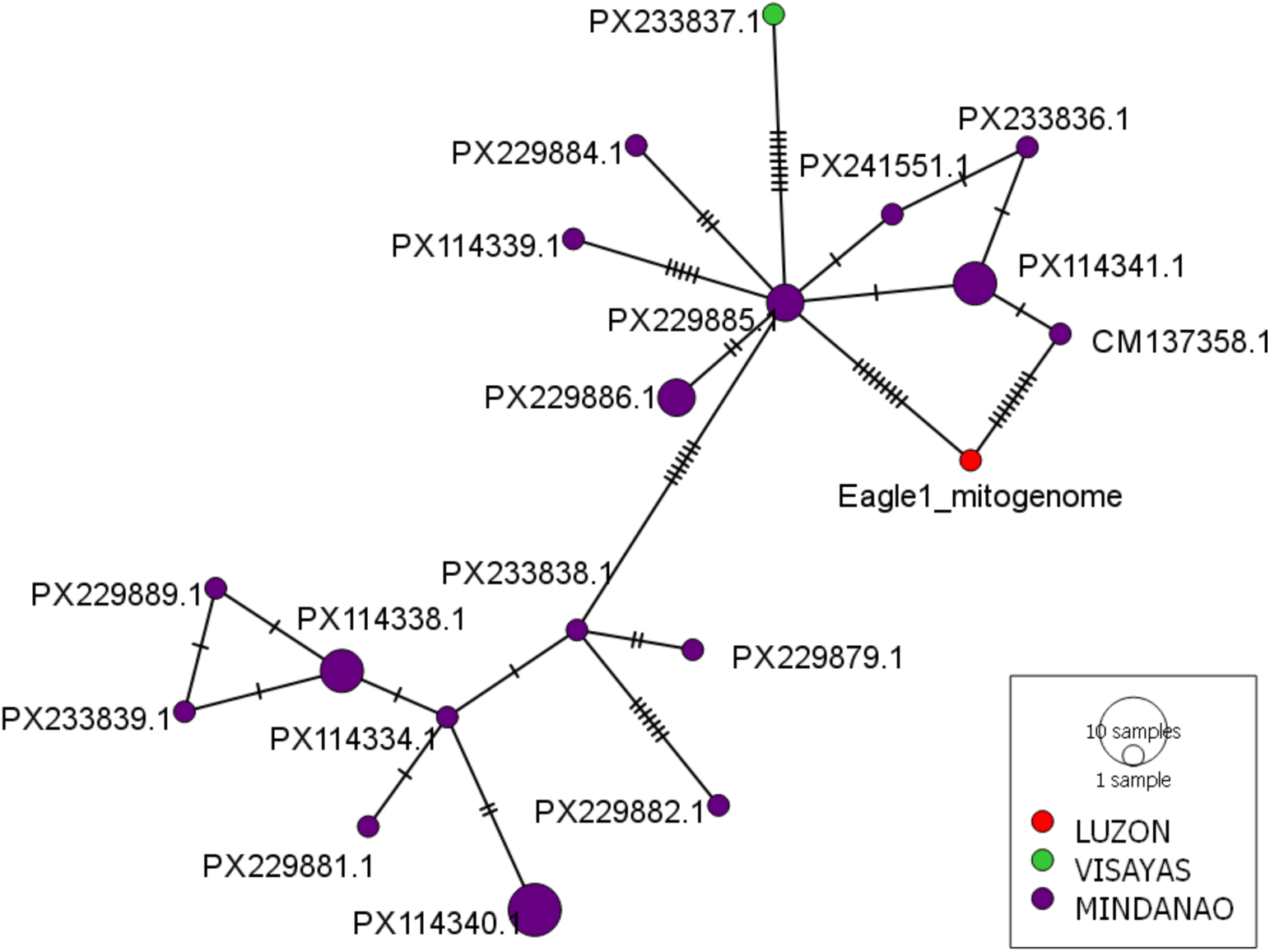
Mitochondrial Network Analysis of Philippine Eagles found throughout the Archipelago

## DISCUSSION

To broaden the geographic scope of genetic information for the Philippine Eagle (*Pithecophaga jefferyi*) and to generate a reference genome for this critically endangered species, we sequenced the genome of a female juvenile named “Binuang” from the northern island of Luzon. Previous genetic studies were limited primarily to individual birds from the southern island of Mindanao (Luczon et al., 2014; Ong et al., 2019; Bacus et al., 2025). Our reference genome, which we have named UST_PhEagle 1.0, represents a high-quality, near-complete avian assembly with a well-resolved gene annotation. It comprises 1.346 Gb with superior contiguity (contig N50 = 45.4 Mb; L50 = 4) and no detectable assembly gaps. Genome completeness is near complete (BUSCO aves_odb12 = 99.92%), with minimal duplication (0.21%) and negligible missing content (0.02%). A consensus mitogenome sequence of 19,377 bp was also generated.

The compact genome size (1.35 Gb) and low repeat content (15.78%) are consistent with established avian genomic architecture, where the typical avian genome ranges from 1 Gb to 2.1 Gb, with a lower repeat content, from 5% to 31% of the total genome, as compared to mammals, from 35% to 52% (Wu et al., 2021). The number of genes in our annotation (23,847 protein-coding genes) lies at the upper-end of the typical range (17,000 to 22,000 genes) for avian genomes (Zhang et al., 2014). While the total gene count is higher than typically reported for avian genomes, exon–intron structure and transcript-to-gene ratios are consistent with known avian gene architecture, suggesting that the inflation likely reflects conservative gene-model prediction rather than assembly fragmentation.

The proportion of genes assigned to orthogroups in the Philippine eagle genome (91.1%) is consistent with values reported for other avian genomes, including the Harpy eagle and Zebra finch (∼90–95%), and slightly lower than the chicken genome (∼92–96%) (International Chicken Genome Sequencing Consortium, 2004; Warren et al., 2010; Zhang et al., 2014; Canesin et al., 2024). Similarly, the fraction of unassigned genes (8.9%) falls within the expected range for birds (∼5–10%). The proportions of species-specific genes (∼3%) and orthogroups are comparable to those of other avian taxa, including raptors and passerines, and do not indicate gene-model inflation (Zhang et al., 2014). MitoZ annotation revealed that the consensus mitogenome sequence is circular and contains the complete set of 13 protein-coding genes, 22 tRNA genes, and 2 rRNA genes. Together, these results support the completeness and reliability of the annotation and suggest typical levels of lineage-specific gene content for an island-endemic raptor genome.

Avian genomes are known for their low repeat content compared with those of other animals (Zhang et al., 2014; Kapusta & Suh, 2017; Galbraith et al., 202; Griffin et al., 2024). Our Philippine eagle genome exhibits a moderately elevated transposable element (TE) content (∼15.8%), similar to the TE content (16.33%) found in the Harpy Eagle (*Harpia harypyja*) genome (Canesin et al., 2024), both of which are higher that the typical avian range of ∼4-10% (Zhang et al., 2014). This elevated repeat fraction likely reflects lineage-specific expansion of transposable elements, particularly CR1 LINEs, which dominate avian repeat landscapes (Kapusta & Suh, 2017). Similar increases in repeat content have been documented in other non-passerine birds, including raptors, suggesting that moderate repeat enrichment is not uncommon in these groups.

We calculated a genome-wide heterozygosity (H) of 0.020% for *Pithecophaga jefferyi*, indicating extremely low levels of genetic variation relative to other avian species (Table 3). In comparison, IUCN Red List species of least concern, such as the budgerigar (*Melopsittacus undulatus*) and little egret (*Egretta garzetta*), exhibit substantially higher genome-wide heterozygosity (0.25–0.43%), with an average of ∼0.249% across 32 species (Li et al., 2014). Even among large raptors, which typically display reduced genetic diversity, heterozygosity values remain higher than those observed in the Philippine eagle, with estimates of ∼0.04% reported for both the bald eagle (*Haliaeetus leucocephalus*) and the white-tailed eagle (*Haliaeetus albicilla*) (Li et al., 2014). As such, the Philippine eagle falls at the lower extreme of the avian heterozygosity spectrum and is comparable to, or below, values reported for IUCN Red List endangered taxa such as the crested ibis (*Nipponia nippon*), which has a whole-genome heterozygosity (H) of 0.04% (Li et al., 2014). Our measure of a low level of genetic variation determined from a whole-genome calculation (H) coincides with the finding that the mean nucleotide diversity (π) for the mitochondrial genome of *Pithecophaga jefferyi* is also very low at 0.054% (Bacus et al., 2025). Both estimates of genetic variation indicate a reduced effective population size (*Ne*) for the Philippine eagle and demographic contraction over time.

We found strong synteny between the genomes of our Philippine Eagle (*Pithecophaga jefferyi*) and the Harpy Eagle (*Harpia harypyja*), which has one of the best reference genomes of the *Accipitriformes* (Canesin et al., 2024). The synteny illustrates the autosomal stability contrasted with the sex-chromosomal plasticity commonly found in avian genomes: Birds are known for having highly syntenic genomes (Zhang et al., 2020; Feng et al., 2020). They even share large syntenic blocks with turtles, suggesting that bird genomes retain their ancestral reptilian roots (Griffin et al., 2024; Liu et al., 2025). However, we also noted the well-documented rapid structural evolution observed in the W and Z chromosomes. Sex chromosomes often undergo suppressed recombination and accumulate structural variants, such as inversions, much faster than autosomes (Sigeman et al., 2021; Sigeman et al., 2024).

The phylogenetic placement of the Philippine eagle, *Pithecophaga jefferyi*, has historically been contentious, with competing hypotheses based on morphology and limited molecular data alternatively placing the species with harpy eagles, booted eagles, or serpent eagles (Lerner & Mindell 2005; Haring et al. 2007). Early morphological classifications placed *Pithecophaga* within a harpy-eagle assemblage alongside *Harpia*, based on similarities in size, forest ecology, and cranial morphology (Brown & Amadon, 1968). Subsequent phylogenetic and comparative analyses instead emphasized affinities with aquiline and hawk-eagles, particularly Aquila and Spizaetus, and demonstrated that traditional harpy-based groupings reflected morphological convergence rather than shared ancestry (Lerner & Mindell, 2005; Gamauf et al., 2005). Genome-scale phylogenies (Catanach et al., 2024) have since resolved this uncertainty, consistently placing *Pithecophaga* within the serpent-eagle radiation. The topology recovered here is consistent with recent phylogenomic analyses and supports a close affinity between *Pithecophaga* and serpent eagles rather than with the aquiline or harpy eagle lineages.

The PSMC trajectory reveals a dynamic demographic history of the Philippine eagle, with a relative peak around 1 MYA, of about 40,000 individuals. This peak was followed by a gradual but sustained decline toward the present. Between about 1 MYA and 100 KYA, the effective population (*Ne*) gradually decreased from about 40,000 to about 4,000 individuals, a population size that remained stable until 10 KYA. Interestingly, this stability coincides with the submersion of the Sundaland land bridges between 18 KYA and 5 KYA, creating the Philippine archipelago (Kim et al., 2023). Today, the estimated average population is 392 potentially breeding pairs or 784 mature individuals (BirdLife International, 2025), which is only about 20% of the population size ten thousand years ago. For comparison, the PSMC trajectory of the Harpy eagle had a relative peak around 1 MYA, of about 8,000 individuals, followed by periods of increase and decrease in *Ne* until about 20 KYA, when the population of about 4,000 individuals entered into a precipitous decline, reaching an effective size of less than 10% of the peak (Canesin et al., 2024).

There has been speculation that the precipitous decline in the Harpy eagle population may have been linked to human activity (Canesin et al., 2024). From the fossil record, anatomically modern humans were already in the Philippine archipelago about ∼30 KYA (Détroit et al., 2004; Détroit et al., 2013), though there is evidence that other hominin species had inhabited the islands ∼66 KYA (Mijares et al., 2010; Détroit et al., 2013). Thus, the decline in the Philippine eagle population appears to coincide with the appearance of *Homo* in the archipelago, though there is no direct evidence to determine whether the former was caused by the latter.

*In toto,* the demographic trend is consistent with our calculated low genome-wide heterozygosity estimate (0.020%), indicating limited extant neutral genetic variation and supporting a history of prolonged small effective population size, demographic contraction, or recurrent bottlenecks. Our findings confirm the Philippine eagle’s status as a critically endangered species.

Next, minimum spanning network analysis revealed that “Binuang” occupies a terminal position, separated from the central haplotype cluster occupied by eagles from the island of Mindanao, specifically the individual birds from the Eastern Mindanao Biodiversity Corridor (Bacus et al., 2025), by multiple mutational steps. This placement suggests that our Luzon eagle represents a genetically distinct mitochondrial lineage from the Mindanao eagles, likely reflecting the historical isolation of different island populations. As such, our data suggests that there may be at least two conservation Management Units (MUs) for the Philippine eagle. Conservation Management Units are populations with significant divergence of allele frequencies at nuclear or mitochondrial loci, regardless of the phylogenetic distinctiveness of the alleles (Moritz, 1994a; Moritz, 1994b). Strikingly, there is evidence that birds from Luzon possess adaptive traits that distinguish them from their counterparts on other islands of the archipelago (Abano et al., 2016).

If confirmed by genome-wide comparisons, this may mean that each MU should be managed with independent demographic and genetic goals, with Luzon eagles not being replaced or supplemented by Mindanao stock and vice versa. Moreover, it may mean that captive breeding programs for the Philippine eagle should maintain lineage separation to preserve locally adapted gene combinations, ensuring that reintroduced individuals are best adapted to their local environments. Nonetheless, our finding that “Binuang” represents a distinct mitochondrial lineage suggests that the Luzon population of Philippine eagles carries irreplaceable genetic diversity. Its loss would represent an evolutionary dead end with no equivalent in the Mindanao population

Our study has at least one significant limitation: We were unable to perform High-throughput Chromosome Conformation Capture (Hi-C), which would have allowed us to better capture the chromosomal structure of our genome (Belton et al., 2012). Therefore, we were unable to confidently assign the macrochromosomes and minichromosomes of the *Pithecophaga jefferyi* karyotype, which has been described as 2n = 66, 3 pairs of relatively large biarmed autosomes, 13 pairs of acrocentrics, three clearly distinguishable groups of medium-sized to small biarmed elements, and 8 microchromosomes (De Boer and Sinoo, 1984). Nonetheless, we achieved high contiguity with long reads, enabling us to map syntenic relationships between our genome and other reference genomes of the Accipitridae.

Finally, in the future, we will need to obtain additional eagle genomes from Luzon to confirm the minimum spanning network analysis, which suggests that birds from the different island regions of the archipelago constitute distinct genetic lineages. This will be important for future management and conservation of the critically endangered national bird of the Philippines.

## ACKNOWLEDGEMENTS

We thank Michael Bacus (Philippine Genome Center, Mindanao) and Matthew van Dam (California Academy of Sciences) for discussion. Sequencing of the Philippine Eagle genome was funded by a research grant from Plasmidsaurus.com. The image of the Philippine Eagle “Noblest Flyer Philippine Eagle,” displayed in Figure 1A, was taken by Shmlongakit and used here under a CC BY-SA 4.0 license with no modifications. N. Austriaco has been designated a Balik Scientist of the Republic of the Philippines.

